# Gut Mycobiome Dysbiosis is Linked to Hypertriglyceridemia Among Home Dwelling Elderly Danes

**DOI:** 10.1101/2020.04.16.044693

**Authors:** Hajar Fauzan Ahmad, Josue Leonardo Castro Mejia, Lukasz Krych, Bekzod Khakimov, Witold Kot, Rasmus Leidesdorff Bechshøft, Søren Reitelseder, Grith Westergaard Højfeldt, Søren Balling Engelsen, Lars Holm, Dennis Sandris Nielsen

## Abstract

Gut microbial dysbiosis have been linked to frailty in elderly, yet the presence of fungal communities and their possible association with host health are little understood. This study attempts to identify gut microbial fungal associations with the progression of atherogenic dyslipidemia in a population of older adults by investigating the interplay between dietary intake, gut mycobiome composition, plasma and fecal metabolome and anthropometric/body-composition measurements of 99 Danes aged 65 to 81 (69.57 ± 3.64) years. The gut mycobiome composition were determined by high-throughput sequencing of internal transcribed spacer (ITS2) gene amplicons, while the plasma and fecal metabolome was determined by GC-MS. The gut microbiome of the subjects investigated is home to three main eukaryotic phyla, namely Ascomycota, Basidiomycota and Zygomycota, with genera *Penicillium, Candida*, and *Aspergillus* being particularly common. Hypertriglyceridemia was associated with fewer observed fungal species, and Bray-Curtis dissimilarity matrix-based analysis showed significant (*p* < 0.05) clustering according to fasting levels of circulating plasma triglycerides (Tg) and very low-density lipoprotein (VLDL) cholesterol fasting levels, respectively. Higher levels of Tg and VLDL cholesterol significantly associates with increased relative abundance of genus *Penicillium*, and *Saccha:ramyces* likely mediated by a higher dietary fatty acids intake (*p* < 0.05), and *Saccharomyces, Debaryomyces, Candida, Agaricus* and *Starmerella* were moderately associated with SCFAs groups. Collectively, these findings suggest that gut mycobiome dysbiosis on older adults is associated with hypertriglyceridemia, a known risk factor for development of cardiovascular disease.

## 1. Introduction

Age is known as a dominant cardiovascular disease (CVD) risk factor in both older men and women, including other multiple disorders such as atherosclerosis [1], obesity [2], hypertension [3], dyslipidaemia [4], and hypertriglyceridemia [5]. Despite the close association with genetics and other health disorders, the interactions between nutrition and gut microbiome are increasingly recognised for their contribution to CVD development [6], [7], Gut microbiota (GM) dysbiosis has previously been found to be associated with frailty in elderly people as well as being a risk factor for metabolic disorders [8]–[13] and other diseases like cancers [14], [15]. Thus, maintaining a diverse core gut microbiome has been proposed as a possible approach for embracing healthy ageing [16]–[18].

Research on the GM of elderly has primarily focused on the bacterial components, largely ignoring fungi, archaea and viruses [10], [19]. Previous studies have characterized human gut fungal communities from diverse age groups [20]–[23]. The fungal component of the gut microbiome of healthy individuals has been reported to be dominated by *Saccharomyces, Malassezia*, and *Candida* [21], [24], [25]. Moreover, colonisation of opportunistic fungal pathogens in the gut can induce dysregulation of host immune responses thereby influencing the disease prognosis. Recent studies show that fungi have significant effects in the gut milieu despite their small proportion in number as compared to bacteria [26]. Gut mycobiome dysbiosis; which refer to an imbalance microbial community composition, including symbiont loss, pathobiont or opportunist outgrowth, altered inter-microbial competition, and disturbed microbial diversity of the gut mycobiota, has been associated with irritable bowel syndromes [27], autoimmune [28], obesity [22], cancers and carotid atherosclerosis [29]. However, the role of the gut mycobiome in developing hypertriglyceridemia among the ageing population has often been neglected.

To date, the best-known mechanism by which the elevation of triglycerides (Tg) and very low density level (VLDL) cholesterol levels have been associated with subclinical atherosclerosis and dubbed as independent risk factors for CVD [30]. Several large studies suggest that hypertriglyceridemia due to increased Tg levels is related to increased levels of remnant lipoproteins in promoting atherogenesis [31], [32], Previous study shown that high-fat diet feeding result in an increased proportion of lipo-polysaccharide-containing microbiota in the gut [33] and involved in secreting and synthesizing bioactive metabolites that affect the accumulation of postprandial lipoprotein [34], Inevitably, the microbial metabolites are transferred to distant sites through blood circulation system and influence the occurrence of hypertriglyceridemia [35]–[37],

Currently, a high-throughput sequencing approach is becoming important for studying complex microbial community in various ecological setting, and capable to sequence thousands to millions of base pairs in a short period by targeting 18S rRNA, ITS1 or/and ITS2 [38], [39]. The recent effort in improving sequencing methods and databases, it is feasible now to describe the non-culturable fungal populations [40]. Here, we comprehensively explored the gut fungal composition, dietary intake, plasma metabolome, and anthropometric/body-composition measurements among older adult Danes that associated to hypertriglyceridemia. We observed that the fecal mycobiome dysbiosis is strongly associated with elevated of Tg and VLDL cholesterol levels. Collectively, these findings provide a new insight for a noninvasive approach in diagnosis and predicting hypertriglyceridemia, and suggest that manipulation of gut mycobiome communities might be a novel target in the treatment of atherosclerotic CVD in the near future among the elderly.

## 2. Materials and Methods

### 2.1 Study Design and Participants Recruitment

Participants for this study consisted of 99 elderly Danes from the Counteracting Age-related Loss of skeletal Muscle mass (CALM) cohort that recruited in the Greater Copenhagen area through local newspapers, magazines, radio programs, social media, and presentations at senior centers and public events. The details about the inclusion criteria has been described elsewhere [41]. All experiments were performed in accordance with the Declaration of Helsinki II and approved by The Danish Regional Committees of the Capital Region (number H-4-2013-070) and with informed consent from all participants, registered at ClinicalTrials.gov (NCT02034760). All data are protected under Danish Data Protection Agency 2012-58-0004 - BBH-2015-001 I-Suite.

### 2.2 Sample Collection and Processing

After the recruitment, every participant will deliver the fecal samples in an insulated bag with freezing elements to Bispebjerg Hospital, Copenhagen, Denmark, within 24 hours and stored at −60 °C until further analysis. Prior homogenisation, the raw fecal samples were thawed at 4 °C, resuspended in autoclaved Milli-Q water (1:2 feces/water) for 1 min at high speed (Lab Seward, BA7021). The homogenized fecal samples were aliquoted in 2 mL vials for usage in this study. For gut microbiome characterization, 200 mg of the fecal pellet was recovered for DNA extraction using the standard protocol from the PowerSoil^®^ DNA Isolation Kit (MOBIO Laboratories, Carlsbad, CA, USA) supplemented with a bead beating step (FastPrep) to enhance cell lysis. Quality and concentration of isolated DNA was measured using NanoDrop 1000 Spectrophotometer (Thermo-Fisher, DE, USA), and was stored at −⍰20 °C until later use.

### 2.3 The internal Transcribed Spacer 2 (ITS2) Amplification and Sequencing

The gut mycobiome composition was determined using Illumina MiSeq amplicon-based sequencing of ITS2 gene regions with adapters compatible for the Nextera Index Kit^®^ (Illumina, CA, USA). For the library preparation, the primers ITS3_F: 5’-TCG TCG GCA GCG TCA GAT GTG TAT AAG AGA CAG GCA TCG ATG AAG AAC GCA GC −3’ and ITS4_R: 5’-GTC TCG TGG GCT CGG AGA TGT GTA TAA GAG ACA GTC CTC CGC TTA TTG ATA TGC −3’ [38] were used to cover ITS2 regions. While the first polymerase chain reaction (PCR) was performed on a SureCycler 8800 (Agilent Technologies, Santa Clara, USA) using the following temperature profile: denaturation at 95 °C for 5 min; 33 cycles of 95 °C for 20 s, 56 °C for 30 s and 68 °C for 45 s; followed by final elongation at 68 °C for 5 min, the barcoding was performed at 98 °C for 1 min; 12 cycles of 98 °C for 10 s, 55 °C for 20 s and 72 °C for 20 s; elongation at 72 °C for 5 min during the second step of PCR. Amplicon concentrations was determined using Qubit^®^ dsDNA BR Assay Kit (Life Technologies, CA, USA) using a Varioskan Flash Multimode Reader (Thermo Fischer Scientific, MA, USA) at 485/530 nm. Samples were pooled in equimolar concentrations and sequenced on a MiSeq platform (Illumina, CA, USA) using the V3, 2×250bp MID pair-ended kit chemistry.

### 2.4 Analysis of High-throughput Amplicon Sequencing

The raw paired-end reads of ITS2 amplicons data were adapter-trimmed and overlapped using fastp v0.21 [42], Forward and reverse primer sequences at the 5’ and 3’ ends of the merged reads were removed, respectively, using cutadapt v1.18 [43]. The merged and primer-trimmed reads were denoised with dada2 [44] within the QIIME2 v.2021.4 [45]. The ITS2 sequences were searched against the NCBI Fungal ITS database [40] followed by consensus-based classification using the QIIME2 v2021.4 classify-consensus-blast pipeline [45]. Both ASV table and taxonomic classification table were exported using QIIME2 tools into tab-separated values (.tsv format) and manually formatted to generate MicrobiomeAnalyst-compatible input [46] with minor modifications [39],

### 2.4 Clinical Parameters, and Metabolome Data

Phenotypic and blood clinical parameters, short-chain fatty acids (SCFAs), 3-days weighted dietary records have been reported previously [47], These data were used to associate with the gut mycobiome component in the present study.

### 2.5 Bioinformatics and Statistical Analysis

The amplicon data was statistically analyzed using the Marker-gene Data Profiling (MDP) module in MicrobiomeAnalyst [46], [48]. In brief, the Amplicon Sequence Variant (ASV) table, taxonomy table, and metadata were uploaded to the server. The features were filtered using a 20% prevalence mean and 10% variance based on the interquartile range before being normalized using cumulative sum scaling (CSS) [49]. Alpha diversity was measured based on the Chao1 and observed species number metrics with a T-test statistical test, while beta diversity was calculated using analysis of similarities (ANOSIM) based Bray-Curtis distance index, with *p*< 0.05 deemed significant. Following that, EVenn was used to construct Venn diagrams and networks of the core microbiome to depict the shared core mycobiome amongst groups [50].

For multivariate analysis, the correlation between the relative abundance of fungi at genus level and variables ([mycobiome and macronutrients], and [mycobiome and metabolites]) on data (hypertriglyceridemia and normal Tg levels) was predicted using Principle Component Analysis (PCA) biplot and correlation heatmap from *factoextra* version 1.0.7 (https://github.com/kassambara/factoextra/) and *GGally* version 2.1.2 (https://github.com/ggobi/ggally) from, respectively with minor modifications [51]. The significance of Pearson correlation coefficients greater than 0.5 was analyzed by using a two-sided Pearson correlation test with cor.test function at a significance level of 0.05. The significant analyses for macronutrients and metabolites datasets were selected to visualize their distribution between two groups of individuals using boxplots from the *ggplot* package. All the above statistical analysis was performed using R statistical software version 4.2.0. A two-tailed one-sample T-test was carried out to examine the significance of macronutrients and metabolites towards the study group *via* GraphPad Prism version 9.4.1 (GraphPad Software, San Diego, California USA). For all statistical tests, unless stated otherwise, a value of *p* < 0.05 was considered as statistically significant.

## 3. Results

### 3.1 Clinical Characteristics of Healthy Older Danish

In this study, a total of 99 home-dwelling rather sedentary elderly Danes above the age of 65 years without any known diseases were enrolled in the CALM study [41]. At baseline, the blood parameters and anthropometric measurements were determined in order to access the trajectory in healthy ageing among older persons. Generally, all the participants had no systemic disease, did not receive any treatment with drugs that affected glucose and lipid metabolisms, nor did they take antibiotics [41], In this study we stratified the participants according to a newly proposed cut-off of fasting Tg levels; Tg > 1.70 mmol/l among the elderly [52], [53] defining a group of blood plasma hypertriglyceridemia (Hypertriglyceridemia, N = 30) and a normotriglyceridemia (Normal Tg, N = 69). Here, Hypertriglyceridemiagroup displayed the typical features of these phenotypes in comparison with NG group, such as significantly higher BMI (*p* = 0.003), higher blood pressure; diastolic (*p* = 0.05), higher lipid profiles; total cholesterol (*p* = 0.001),HDL (p < 0.001), LDL (*p* = 0.02), and VLDL (*p* = 0.001), and glucose metabolism; OGTT (*p* = 0.009), Hemoglobin A1c (*p* = 0.021), and Proinsulin C-peptide (*p* < 0.001) when compared using Welch t-test. Nevertheless, age and fasting glucose did not present significant differences between the Hypertriglyceridemia and Normal Tg groups (Table 1).

**Table 1.** Demographics of the study participants.

| Features | Overall | Normal Tg Group | Hypertriglyceridemia Group | $p$-value |
| --- | --- | --- | --- | --- |
| Sample size | N = 99 | N = 69 | N = 30 | - |
| Age (years) | 69.57 ± 3.64 | 69.27 ± 3.48 | 70.27 ± 3.94 | 0.106 |
| BMI (kg/cm<sup>3</sup>) | 25.43 ± 3.45 | 24.81 ± 3.29 | 26.87 ± 3.43 | 0.003* |
| **Blood pressure** |  |  |  |  |
| Systolic (mmHg) | 143.37 ± 19.74 | 142.86 ± 21.37 | 144.57 ± 15.54 | 0.347 |
| Diastolic (mmHg) | 84.95 ± 10.74 | 83.79 ± 10.03 | 87.67 ± 11.97 | 0.050* |
| **Blood Lipid profiles (mmol/L)** |  |  |  |  |
| Total cholesterol | 5.72 ± 0.93 | 5.54 ± 0.89 | 6.14 ± 0.91 | 0.001* |
| HDL-cholesterol | 1.80 ± 0.49 | 1.92 ± 0.46 | 1.50 ± 0.43 | <0.001* |
| LDL-cholesterol | 3.23 ± 0.90 | 3.12 ± 0.86 | 3.53 ± 0.96 | 0.020* |
| VLDL-cholesterol | 0.66 ± 0.30 | 0.51 ± 0.14 | 1.04 ± 0.24 | <0.001* |
| Fasting Tg | 1.50 ± 0.76 | 1.11 ± 0.30 | 2.43 ± 0.72 | 0.080 |
| **Blood Glucose Profile (mmol/L)** |  |  |  |  |
| Fasting glucose | 5.41 ± 0.51 | 5.37 ± 0.43 | 5.51 ± 0.59 | 0.115 |
| OGTT 120 glucose | 6.75 ± 1.63 | 6.50 ± 1.60 | 7.35 ± 1.57 | 0.009* |
| Haemoglobin A1c | 35.6 ± 3.15 | 35.19 ± 3.21 | 36.57 ± 2.81 | 0.021* |
| Proinsulin C-peptide (pmol/L) | 707 ± 278 | 623.27 ± 213 | 916.46 ± 314 | <0.001* |

Notes \* $p$ -values are from Welch t-tests for continuous variables, between two groups of Tg levels. Tab $p$ - values < 0.05 considered significant. Abbreviations; BMI – Body Mass Index, HDL – High Density Lipoprotein, LDL – Low Density Lipoprotein, VLDL – Very Low Density Lipoprotein, Tg – Triglycerides, and OGTT – Oral Glucose Tolerance Test

### 3.2 Fungal Community Composition and Diversity in Hypertriglyceridemia and Normal Tg Groups

Overall, a total of 99 fastq files were generated from Illumina Miseq sequencing and obtained 7,830,381 high-quality ITS2 sequence reads, with an average of 79,094 reads per sample (min = 2832, max = 481718). The mapped sequence reads yielded 1712 ASVs belonging to 3 phyla, 16 classes, 35 orders, 54 families, 82 genera, and 104 fungal species. It is important to note that the sample size for each analysis may vary owing to missing data in certain parameters (VLDL levels, macronutrients, metabolites), as a consequence of individuals dropping out of the study.

The taxonomy bar plots revealed that *Ascomycota* dominated both hypertriglyceridemia (n=30) and normal Tg (n= 69) groups, accounting for 96% and 99% of total ASVs, respectively (Figure 1A). Both groupings mainly consisted of *Penicillium, Saccharomyces, Geotrichum*, and unclassified *Ascomycota* at the genus level (Figure 1B). *Penicillium* had the highest proportion in hypertriglyceridemia, contributing to 54.7% of total fungi abundance, followed by 20.7% of *Saccharomyces*. Meanwhile, the normal Tg group included more *Saccharomyces* (31.7%) and *Geotrichum* genera (30%), accompanied by various fungal genera in minor abundance. Additional information regarding fungi composition can different taxonomy levels can be found in the supplementary section (Figure S1, Table S1).

**Figure 1:**
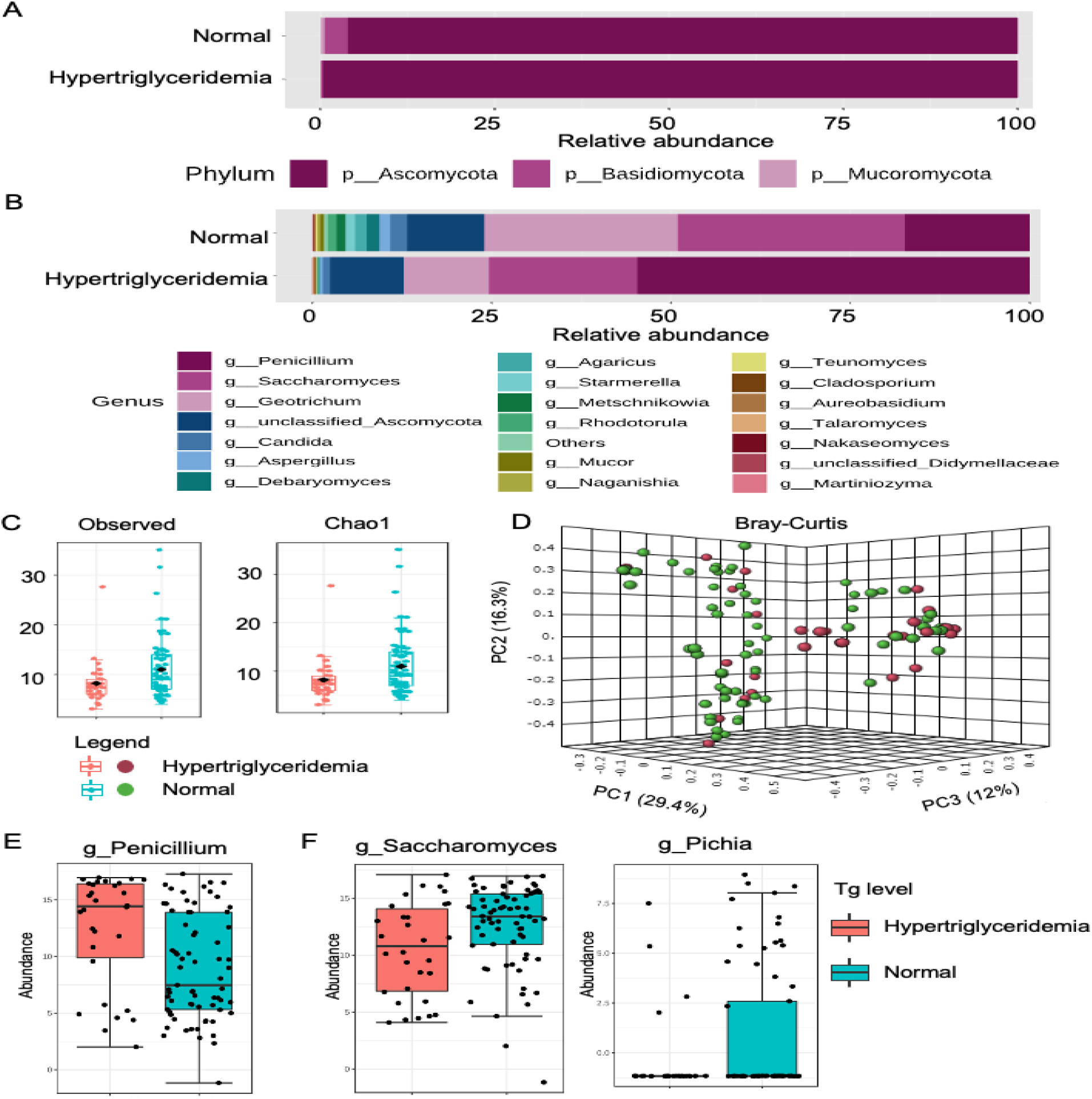
Profiling of gut mycobiome linked with Tg levels using ITS 2 gene region. The identified taxa at the A) phy-lum; and B) genus levels were expressed as percentage abundance in merged samples. Only the top 20 genera are shown at the genus level, with the remainder classified in Others. C) Alpha diversity analysis using observed species number and Chaol indices revealed significant differences in species richness at the genus level between two groups (p-value= 0.0123). D) Beta diversity analysis based on Bray-Curtis dissimilarity metric reveals significant variation between two groups (R = 0.010, p-value < 0.008). The boxplots depicted the abundance of significant fungal taxa prevalent in E) hypertriglyceridemia; and F) normal Tg levels using metagenomeSeq. The red and blue boxplots repre-sented hypertriglyceridemia and normal Tg level, respectively.

The alpha and beta diversity measured the fungal diversity within and between the communities, respectively. The richness of fungi communities was significantly higher in normal Tg than in hypertriglyceridemia, as evidenced by observed species number and Chao1 estimator in alpha diversity analysis (both p=0.0123, T-test= −2.5671; Figure 1C). In addition, the ANOSIM-based Bray-Curtis distance index in beta diversity analysis indicated significant differences in fungal population between hypertriglyceridemia and normal Tg groups (*p*< 0.008, [ANOSIM]R=0.1049). The differences between the fungal communities in two different groups were illustrated in the principal coordinate analysis (PCoA) 3D plot (Figure 1D). On the other hand, the clustering analysis highlighted the distribution pattern of the fungal genus based on the Pearson correlation coefficient. According to the bar plot, *Pichia, Kurtzmaniellahia, Cophinforma, Tausonia, Clavispora, Hanseniaspora, Saccharomyces, Teunomyces*, and *Agaricus* genera found abundant in normal Tg levels, whereas *Penicillium* was moderately correlated with hypertriglyceridemia (Figure S3). The differential abundance analysis of the metagenomeSeq model with zero-inflated Gaussian distributions revealed that *Penicillium*was significantly prevalent in hypertriglyceridemia (p-value=0.001, FDR= 0.006314; Figure 1E), whereas *Pichia* (p-value=1.68e^-12^, FDR=2.65e’^-11^) and *Saccharomyces* (p-value=0.005, FDR=0.012) were enriched in normal Tg levels (Figure 1F). The complete metagenomeSeq analysis can be found in Table S3, with a False Discovery Rate (FDR) adjusted p-value < 0.05 considered significant.

### 3.3 Fungal community composition and diversity in high and normal VLDL groups

The taxonomy distribution among high (n=29) and normal VLDL (n=67) groups were relatively similar to hypertriglyceridemia and normal Tg levels groups. The fungal composition was predominated by *Ascomycota* phyla in both groups (High VLDL=99.6%, normal VLDL=96.5%), represented by *Penicillium, Saccharomyces, Geotrichum*, and unclassified *Ascomycota* genera. Of these, the highest proportion of *Penicillium* genera was detected in the high VLDL group (53.2%), whereas the normal VLDL group constituted a larger proportion of *Saccharomyces* (32.5%) and *Geotrichum* (28%) genera. Unclassified *Ascomycota* had a comparable prevalence in both groups, contributing to 9.9% to 11% of overall abundance in the normal and high VLDL groups, respectively. Figure S2 and Table S2 provide additional information on fungi composition based on VLDL levels.

Likewise, the observed species number and Chao1 metrics reflected significant differences in fungal diversity between the normal and high VLDL groups, with normal VLDL exhibiting greater variation than high VLDL (both p=0.0183, T-test=2.4181; Figure 2C). Beta diversity using the ANOSIM approach showcased distinct fungi variation at the genus level across two groups, as evidenced by the Bray-Curtis distance index with *p*-value < 0.003 and [ANOSIM]R= 0.11841 (Figure 2D). Besides, 14 out of 25 genera was found to associated with normal normal VLDL levels (Figure S3). Figure 1E illustrated the significant prevalence of *Saccharomyces* (p-value=0.003, FDR=0.018), *Pichia* (p-value=l.07e^-08^, FDR=1.07e^-07^) and *Teunomyces* genera (p-value=0.0002, FDR=0.002) based on metagenomeSeq analysis (Table S4).

**Figure 2:**
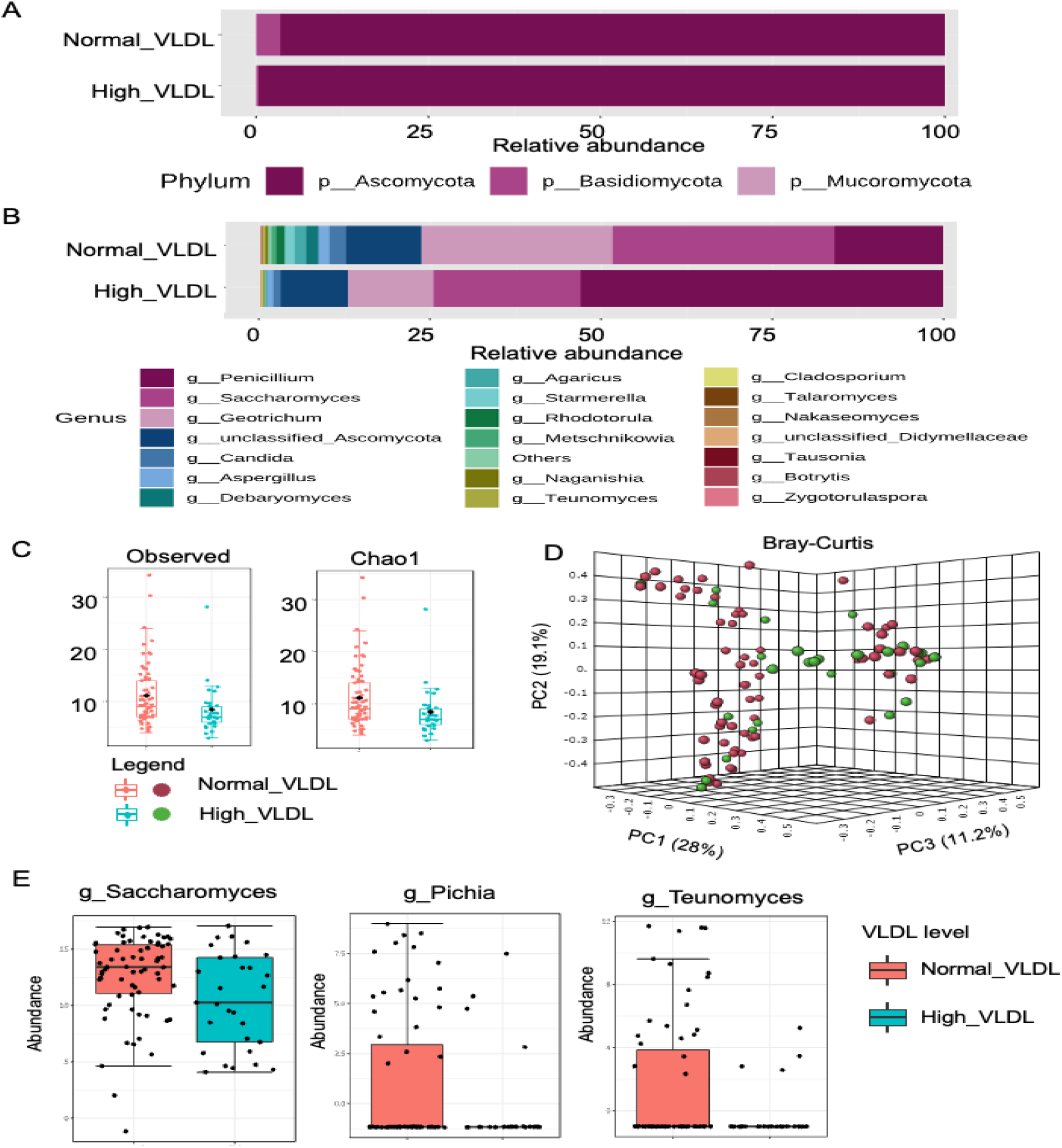
Profiling of gut mycobiome linked with VLDL levels using ITS 2 gene region. The identified taxa at the A) phylum; and B) genus levels were expressed as percentage abundance in merged samples. Only the top 20 genera are shown at the genus level, with the remainder classified in Others. C) Alpha diversity analysis using observed species number and Chaol indices revealed significant differences in species richness at the genus level between two groups (p-value = 0.0183). D) Beta diversity analysis based on Bray-Curtis dissimilarity metric reveals significant variation between two groups (R = 0.012, p-value < 0.003). E) The boxplots depicted the abundance of significant fungal taxa prevalent in normal VLDL level using metagenomeSeq. The red and blue boxplots represented normal VLDL and high VLDL, respectively.

### 3.4 Core mycobiome at Tg and VLDL Levels Reveals the Interconnectedness

The fungal genera that were consistently present across the sample groups were identified at a 20% sample prevalence. *Penicillium, Saccharomyces*, unclassified *Ascomycota*, and *Geotrichium* genera were common in the core mycobiome detected at Tg and VLDL levels. Hypertriglyceridemia had the most prevalence of *Penicillium* (preva-lence=0.8), followed *by Saccharomyces*, unclassified *Ascomycota*, and *Geotrichium*.Meanwhile, *Saccharomyces* (prevalence = 0.8) was abundant among normal Tg groups, followed by unclassified *Ascomycota, Penicillium*, and *Geotrichum*. The core mycobiome based on VLDL levels was similar to Tg levels, with high VLDL matching with hypertriglyceridemia and normal VLDL matching with normal Tg groups (Figure S3).

A Venn network was constructed to demonstrate the connection of the core mycobiome at the genus level between Tg levels and VLDL levels (Figure 3A). A total of 16 fungi were found in groups with normal Tg and normal VLDL levels, whereas 4 fungi were common in hypertriglyceridemia with high VLDL groups. Interestingly, 12 fungi were interconnected among four groups, including *Penicillium, Saccharomyces*, unclassified *Ascomycota*, and *Geotrichium* genera. *Cladosporium* was detected in all three groups, except hypertriglyceridemia, while *Tausonia* was only found in the normal Tg group. Besides, a Venn diagram summarized the number of core mycobiome of four groups, with normal Tg and VLDL having more fungi genera (30 and 29 fungi, respectively) than hypertriglyceridemia and high VLDL (16 and 17 fungi, respectively) (Figure 3B). The complete output of the Venn diagram is available in the supplementary section (Table S5).

**Figure 3:**
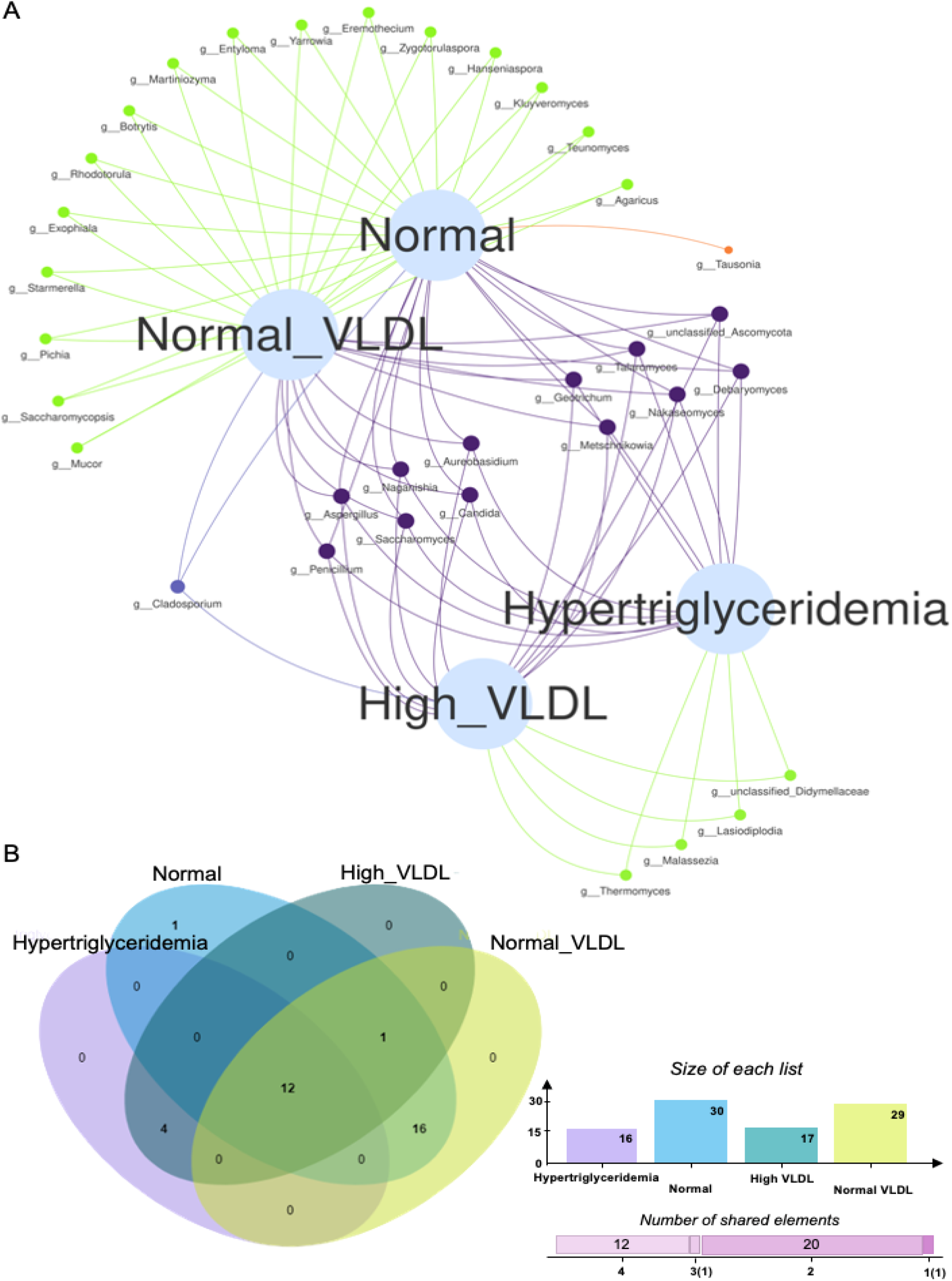
Venn diagram-based analysis revealed the association of core mycobiota between Tg and VLDL levels. A) Networks of core mycobiota shared by Tg and VLDL groups. Hyperglyceridemia shared connection with high VLDL, while mycobiota in normal Tg levels is associated with normal VLDL. B) Venn diagram of core mycobiota composition. Hypertriglyceridemia denoted in purple, Normal Tg levels in blue, High VLDL in grey and Normal VLDL in yellow. The number represents the number of core mycobiota belong to each group. The numbers shown in overlapping regions represent the number of shared fungi. Notice that 16 fungi genera are shared between normal Tg and VLDL levels, meanwhile 12 fungi genera are commonly shared with all groups.

### 3.5 Correlation Analysis of Fungal Communities with Macronutrients among Hypertriglyceridemia Individuals

The association between macronutrients and fungal communities in hypertriglyceridemia and normal Tg groups was analyzed using PCA analysis. The first two principal coordinates (Dim1= 19.7%, Dim2= 11.1%) described the variability in macronutrients and mycobiome between the hypertriglyceridemia and normal Tg groups. As illustrated in Figure 4A, the macronutrients, particularly polyunsaturated fatty acids, monounsaturated fatty acids, fat, protein, sugars, carbohydrate available, saturated fatty acids, and dietary fiber, were grouped together and intimately linked. In the meantime, legumes, vegetable oil, alcohol, butter, and other fat, as well as fungal communities, were isolated from the clustered groupings and weakly correlated with one another. Significant clustering of hypertriglyceridemia and normal Tg groups was observed, with ellipses overlapping between the two groups. Hypertriglyceridemia groups were distributed around fatty acids groupings, while normal Tg groups encompassed a broader range of macronutrients. The correlation heatmap depicted the degree of Pearson correlation coefficient between macronutrients and mycobiome, revealing a significant strong positive correlation between macronutrients (p< 0.05; Figure 4B, Table 1). While legumes were strongly correlated to the genus *Agarícus*, other dietary elements such as dietary fiber and fatty acids have no significant association with mycobiome profiles.

**Figure 4:**
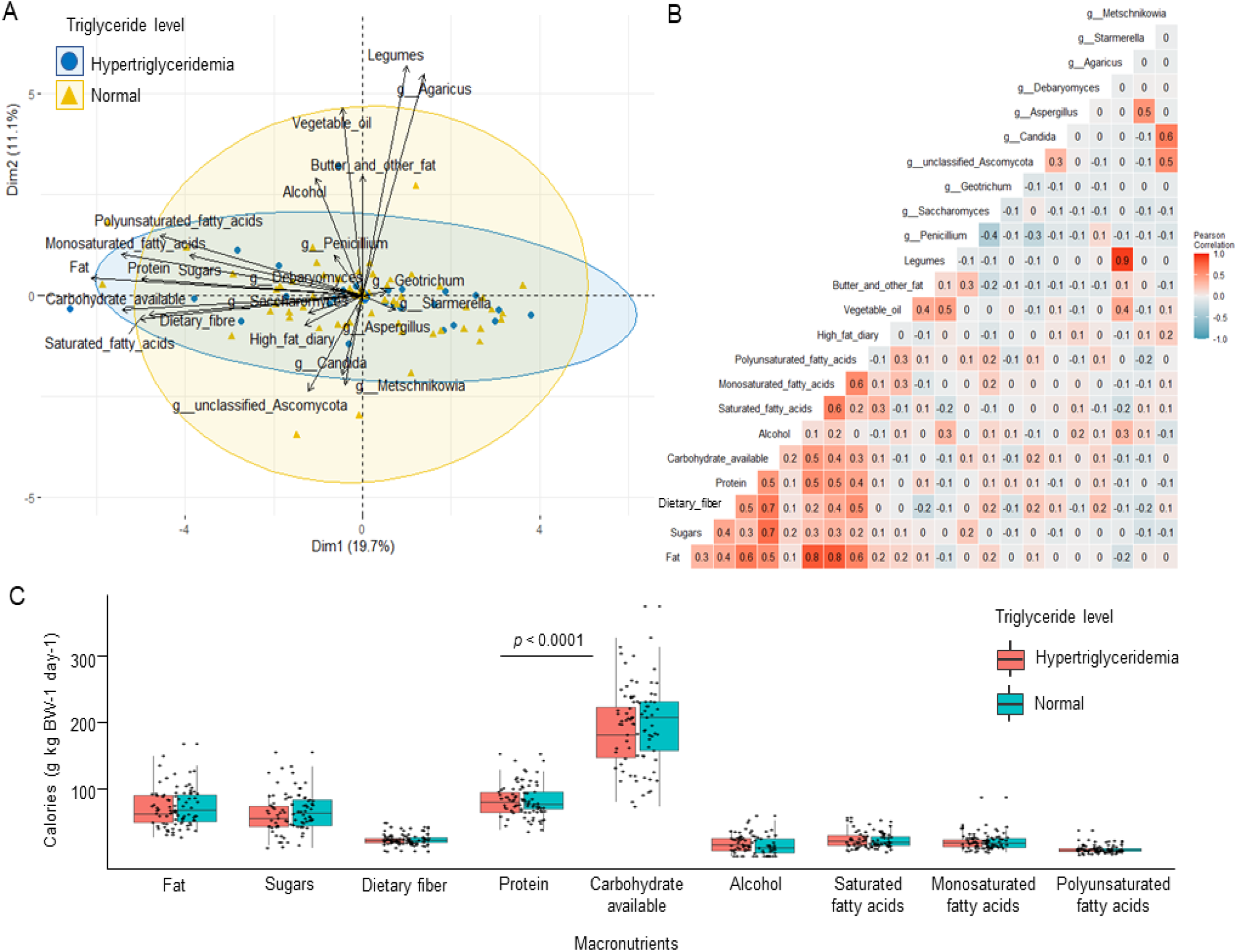
Multivariate statistical analysis of dietary nutrients with the top ten fungi genera. A) Biplots of principal component analysis (PCA) demonstrated the explanatory variables as vectors (black lines) and points. Blue circles represent hypertriglyceridemia and yellow triangles, normal Tg levels. The ellipses indicated the grouping. Positively correlated variables have vectors pointing in the same direction, while negatively correlated variables have vectors pointing in opposite directions. B) The strength of association between fungi genera and variables was expressed as Pearson correlation coefficient in correlation matrix, represented by the colour strength and the numerical value. Positive correlation is shown by red while negative correlation is denoted by blue. C) Boxplots described the type of macronutrients linked with the of distribution of calories (Cal kg body weight^-1^ day^-1^). One-sample t-test determined that each macronutrient has a different population mean (*p* < 0.05).

To further explore the effect of macronutrients among hypertriglyceridemia and normal Tg groups, the significant macronutrients were chosen to visualize the contribution of macronutrients in calories (Cal kg body weight^-1^ day^-1^) among the two groups. The high calories intake indicated the high energy intake, with carbohydrate available (l95±6.479) ranking first, followed by protein (82.04±2.539), fat (72.97±3.096), sugars (64.73±3.085), dietary fiber (24.43±0.857), saturated fatty acids (23.55±l.2l4), mono-saturated fatty acids (2l.42±l.29l), alcohol (l6.48±l.544), and polyunsaturated fatty acids (10.l5±0.624) (Figure 4C; Table S6). All the macronutrients tested showed significant differences with p< 0.0001.

### 3.6 Correlation Analysis of Fungal Communities with SCFAs among Hypertriglyceridemia Individuals

The association between targetted metabolites and fungal communities in hypertriglyceridemia and normal Tg groups as depicted in Figure 5A, with all SCFAs (acetic acid, propionic acid, isobutyric acid, butyric acid, isovaleric acid, and valeric acid) grouped and closely related together. The mycobiome was extensively scattered and was unlikely to connect with the targetted metabolites. Likewise, the two groups overlapped in that they both covered *Penicillium, Saccharomyces*, and *Geotrichum*. Despite that the correlation matrix heatmap indicated a strong positive correlation among SCFAs and mycobiome, but no obvious association was identified between them (p< 0.05; Figure 5B, Table 2).

**Figure 5:**
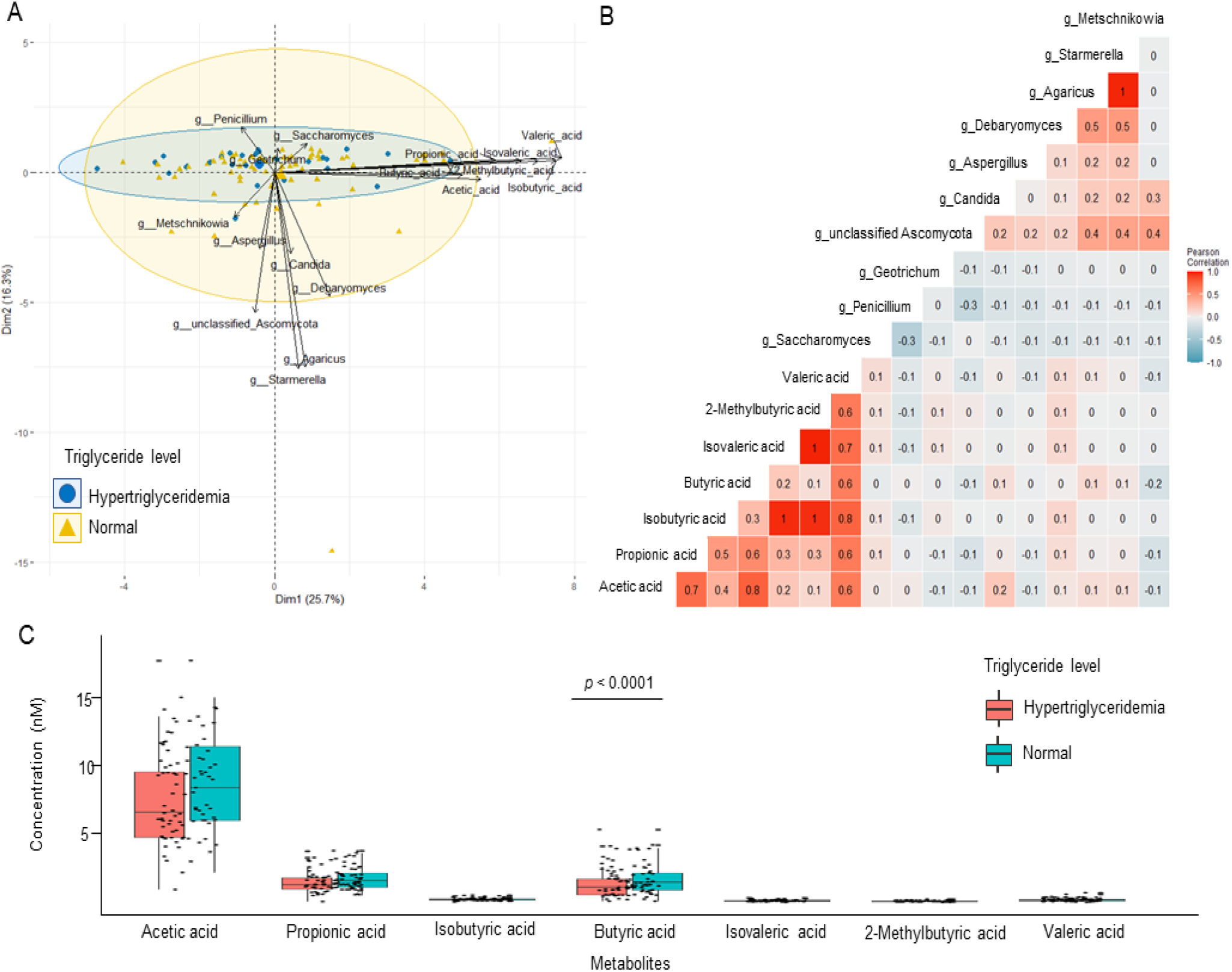
Multivariate statistical analysis of metabolites with the top ten fungi genera. A) Biplots of principal component analysis (PCA) demonstrated the explanatory variables as vectors (black lines) and points. Blue circles represent hypertriglyceridemia and yellow triangles, normal Tg levels. The ellipses indicated the grouping. Positively correlated variables have vectors pointing in the same direction, while negatively correlated variables have vectors pointing in opposite directions. B) The strength of association between targetted metabolites and fungi genera was expressed as Pearson correlation coefficient in correlation matrix, represented by the colour strength and the numerical value. Positive correlation is shown by red while negative correlation is denoted by blue. C) Boxplots depicted the concentration of SCFAs among hypertriglyceridemia and normal TG individuals. One-sample t-test determined that each SCFAs has a different population mean (*p* < 0.05).

**Table 2.** Pairwise comparison of macronutrients and mycobiome with a Pearson correlation coefficient greater than 0.5.

| Variables1 | Variables2 | Correlation coefficient (r >0.5) | Pearson (p<0.05) |
| --- | --- | --- | --- |
| Fat | Protein | 0.6 | 3.31E-11 |
|  | Saturated fatty acids | 0.8 | 1.86E-20 |
|  | Monosaturated fatty acids | 0.8 | 1.08E-19 |
|  | Polyunsaturated fatty acids | 0.6 | 3.14E-09 |
| Sugars | Carbohydrate available | 0.7 | 4.93E-14 |
|  | Carbohydrate available | 0.7 | 2.33E-11 |
| Saturated fatty acids | Monosaturated fatty acids | 0.6 | 8.22E-10 |
| Monosaturated fatty acids | Polyunsaturated fatty acids | 0.6 | 2.60E-08 |
| Legumes | *Agaricus* | 0.9 | 2.49E-33 |
| *Candida* | *Metschnikowia* | 0.6 | 1.68E-10 |

**Table 3.** Pairwise comparison of metabolites and mycobiome with a Pearson correlation coefficient greater than 0.5.

| Variables1 | Variables2 | Correlation coefficient (>0.5) | Pearson ($p < 0.05$) |
| --- | --- | --- | --- |
| Acetic acid | Propionic acid | 0.7 | 3.245557e-16 |
|  | Butyric acid | 0.8 | 1.002635e-19 |
| Propionic acid | Isovaleric acid | 0.6 | 0.001434186 |
| Isobutyric acid | Isovaleric acid | 1 | 2.584355e-55 |
|  | 2-Methylbutyric acid | 1 | 6.412354e-52 |
| Isovaleric acid | 2-Methylbutyric acid | 1 | 8.682254e-71 |
| Valeric acid | 2-Methylbutyric acid | 0.6 | 2.375447e-12 |
|  | Isovaleric acid | 0.7 | 9.395219e-16 |
|  | Butyric acid | 0.6 | 1.283421e-11 |
|  | Isobutyric acid | 0.8 | 8.91927e-19 |
|  | Propionic acid | 0.6 | 2.565943e-12 |
|  | Acetic acid | 0.6 | 4.836536e-12 |
| *Agaricus* | *Starmerella* | 1 | 1.564446e-64 |

Furthermore, the concentration of SCFAs on triglyceride levels was also examined. Acetic acid (7.8±0.325) was the most abundant in both groups, followed by butyric acid (1.636±0.106) and propionic acid (1.727±0.081), with normal Tg groups having a higher concentration than hypertriglyceridemia groups (Figure 5C). All the SCFAs exhibited significantly different means, as evidenced by a one-samples T-test with p< 0.0001 (Table S7).

## 4. Discussion

The causes of hypertriglyceridemia among the elderly can be a result of interactions between genetic precursors [54], non-genetic factors such as unhealthy lifestyle [55], diseases related to metabolic syndromes [56], usage of some types of medicine [57] and high-fat diet [58]. Epidemiological studies consistently demonstrate strong associations of plasma Tg levels that causing hypertriglyceridemia, with risk of atherosclerotic CVD [59], [60]. Most fungal species detected in gut mycobiome studies are considered transient components of the community, and putative of environmental origin, in particular influenced by food-borne fungi and life-style [61], [62], together with other factors such as age, gender and geographical setting [16], [20], [63]. However, due to the dearth of information related to gut mycobiome studies, little is known about its relationship with fecal metabolome and other factors such as environmental effects, diet and life style [64] that may lead to hypertriglyceridemia. Here, we present data showing an association between gut mycobiome dysbiosis and hypertriglyceridemia in a homogeneous and well-characterized healthy cohort of older Danish adults.

Collectively, we found that the richness of the gut mycobiome among the studied population was low within individuals with *Saccharomyces* and *Pichia* genera being common among healthier older Danish. Previous study also showed lower alpha diversity of fungi community as compared to the gut bacterial community [21], [65], and decreasing throughout the course of life due to ageing [20], [23], with *Saccharomyces* [66] and *Candida* [67] genera formed gut commensal. In the present study, *Penicillium* was observed among many of the subjects, and rather predominant among high Tg and VLDL groups. In contrast, previous study indicated that *Cladosporium* are associated with total cholesterol and LDL among carotid atherosclerosis in younger Spainish populations [21], A total of 30 of the included participants had Tg levels above the recommended level of 1.7 mmol/L [68]–[71]. Similarly, a similar pattern of good versus unhealthy VLDL cholesterol levels strongly linked to the mycobiome composition was observed. The particular patterns observed for both plasma lipid markers are expected, termed as atherogenic lipid triad in dyslipidaemias [72], [73]. Tg and lipoprotein metabolism is linked as a result of their similar physicochemical characteristics involving two major organs such as intestine and liver [74]. In the intestine, bile acids emulsify fats into smaller particles which allows lipases to breakdown Tg into fatty acids. Fatty acids can then be absorbed and be used as substrates for chylomicron assembly that contributes to postprandial Tg levels. Subsequently, the gut microbiota will generate SCFAs, secondary bile acids and lipopolysaccharides [75] which activate receptors that regulate postprandial chylomicron production, and absorbtion of SCFAs and bile acids into the portal circulation. Meanwhile, the SCFA can act as substrates for *de novo* lipogenesis and contribute to VLDL production in the liver [33].

Next, a host-associated core microbiome based on network analysis was conducted to explore common groups of fungi that were likely to be particularly important for host biological function [76]. Four fungi were found to be exclusively common in hypertiglyceridemia with high VLDL such as *Thermomyces, Malessezia, Lasiodiplodia* and unclassified *Didymellaceae* that associated with lipase production. Mounting study showed that *Thermomyces* [77]–[80], *Malessezia* [81]–[83], and *Lasiodiplodia* [84]–[86] involves during fats and lipids hydrolysis in production of fatty acids. Another interesting observation was 16 fungi were found solely in helthier groups with normal Tg and normal VLDL levels, which are *Mucor, Saccharomycopsis, Pichia, Starmerella, Agaricus, Exophiala, Rhodotorula, Botrytis, Martiniozyma, Entyloma, Yarrowia, Eremothecium, Zygo-torulaspora, Hanseniaspora, Kluyveromyces* and *Teunomyces*. Despite being signitures of healthy gut due to long-term habitual diets and healthy host physiological states [23], [88], some fungi like *Mucor* was reported to be abundant in the gut of non-obese subjects [22], and confer protection from the risk of CVD [29]. Interestingly, 12 fungi were interconnected among groups such as *Penicillium, Saccharomyces*, unclassified *Ascomycota*, and *Geotrichium* genera. This analysis showed that an upsurge in *Penicillium*genus could be associated with hypertriglyceridemia. However, the utility of *Penicillium*as a biomarker in predicting the progression of atherosclerosis by modulating the Tg and VLDL among older adults is still unclear, and therefore, this association warrants further investigation.

With regard to the dietary intake, the individuals from hypertriglyceridemia group pose high calories intake indicated the high energy intake due consumption of higher carbohydrate, protein, fat, sugars, and dietary fiber as previously reported [47] that highly adhered to the recommended intake of carbohydrates and fibres [89]. However, the intake of saturated and unsaturated fatty acid groups is still higher in hypertriglyceridemia. Furthermore, correlation analysis with macronutrients components support and stand out the relevance of these fungal in hypertriglyceridemia. Particularly, in the case of *Penicillium*, and *Saccharomyces* positively correlate with diet rich in butter, sugar, and other fatty acids groups which are common indicators for higher Tg and VLDL cholesterol in circulating serum of hosts, which have been reported to be associated with signatures in coronary atherosclerotic plaques [90], aneurysms of the carotid artery [91]. Interestingly, other fungi like *Agaricus* and *Debaryomyces* are strongly associated with legumes and moderately associated with dietary fibre, respectively. Based on another strand of study assessing the risk of suboptimal intake of macro- and micronutrients, it is apparent that healthy community-dwelling older Danes consumed more saturated fats and alcohol than recommended by official dietary reference values [92], which is common in the Western diet [93], [94],

SCFA have a variety of advantageous effects on the human energy metabolism, including the metabolism of glucose, lipids, and cholesterol in a variety of tissue types [95]. The acetic, butyric and propionic acid are significantly higher among helthier elderly, which consistant with a study reported previously in human [10] and animal model [96]. Here, we observed that acetic acid is the most predominant and moderately associated with *Candida, Debaryomyces, Agaricus* and *Starmerella*. Previous study described acetate and butyrate as fermentation products of the complex carbohydrates such as dietary fibres, and also the main substrate for the synthesis of cholesterol [97], The *Sac-charomyces, Geotrichum* and *Debaryomyces* also were found to be moderately associated with other SCFAs like propionic, butyric, valeric acids. However, no significant correlations between *Penicillium* and *Aspergilllus*, with any of the SCFAs were identified. This suggests that the production of SCFAs may be driven by bacterial activities as previously reported [98].

## 5. Conclusions

To the best of our knowledge, this is the first study to demonstrate that hypertriglyceridemia among elderly is associated with gut mycobiome dysbiosis characterized by overall reduction of the microbial richness and diversity as well as dysbiosis pattern of the gut mycobiome structure compared to those senior citizens with normal levels of circulating plasma triglycerides. These findings also highlight that the everyday diet shapes the gut mycobiome and host metabolome components among the older citizens. However, it remains unknown whether the microbial markers and patterns identified here are also adaptable to changes in life styles and applicable to other cultures in the world.

## Supplementary Materials

The following supporting information can be downloaded at: www.mdpi.com/xxx/s1, Figure S1: title; Table S1: title; Video S1: title.

## Author Contributions

HFA performed laboratory procedures; DSN, LH, SBE, SR, JLC, HFA designed the study; RLB, SR, GWH, LH collected and provided samples as well as analyzed clinical data; BK carried out metabolome analysis; WK carried out sequencing of libraries, HFA, JLC, ŁK, KF, DSN coupled and analyzed the different datasets of the study; HFA and DSN drafted the manuscript. All authors commented on, added paragraphs and approved the last version of this manuscript.

## Institutional Review Board Statement

All experiments were performed in accordance with the Declaration of Helsinki II and approved by The Danish Regional Committees of the Capital Region (number H-4-2013-070) and data protected under Danish Data Protection Agency 2012-58-0004 - BBH-2015-001 1-Suite for study involving humans.

## Informed Consent Statement

Informed consent was obtained from all subjects involved in the study and registered at ClinicalTrials.gov (NCT02034760).

## Data Availability Statement

The raw sequence data of this study were uploaded to EBI’s ENA under accession codes PRJEB34758 and PRJEB34758.

---

**Fig. S1.**
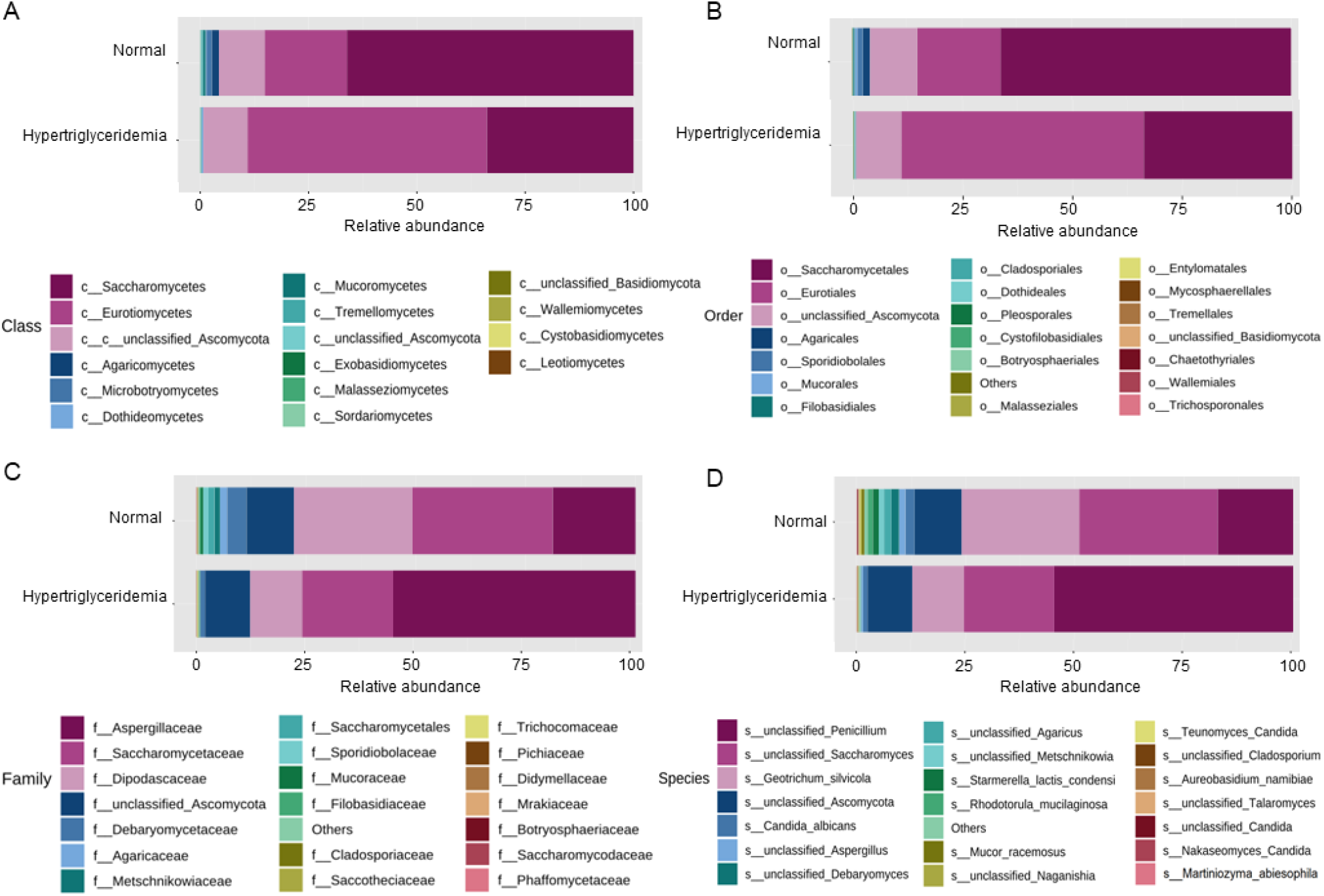
Taxonomy summary of the fungal communities associated with Tg levels. The diagrams depicted the percent-age abundances of fungi at the (A) class; (B) order; (C) family; and (D) species levels in merged samples. Only the top 20 features are shown at the taxonomy level, with the remainder classified in Others.

**Fig. S2.**
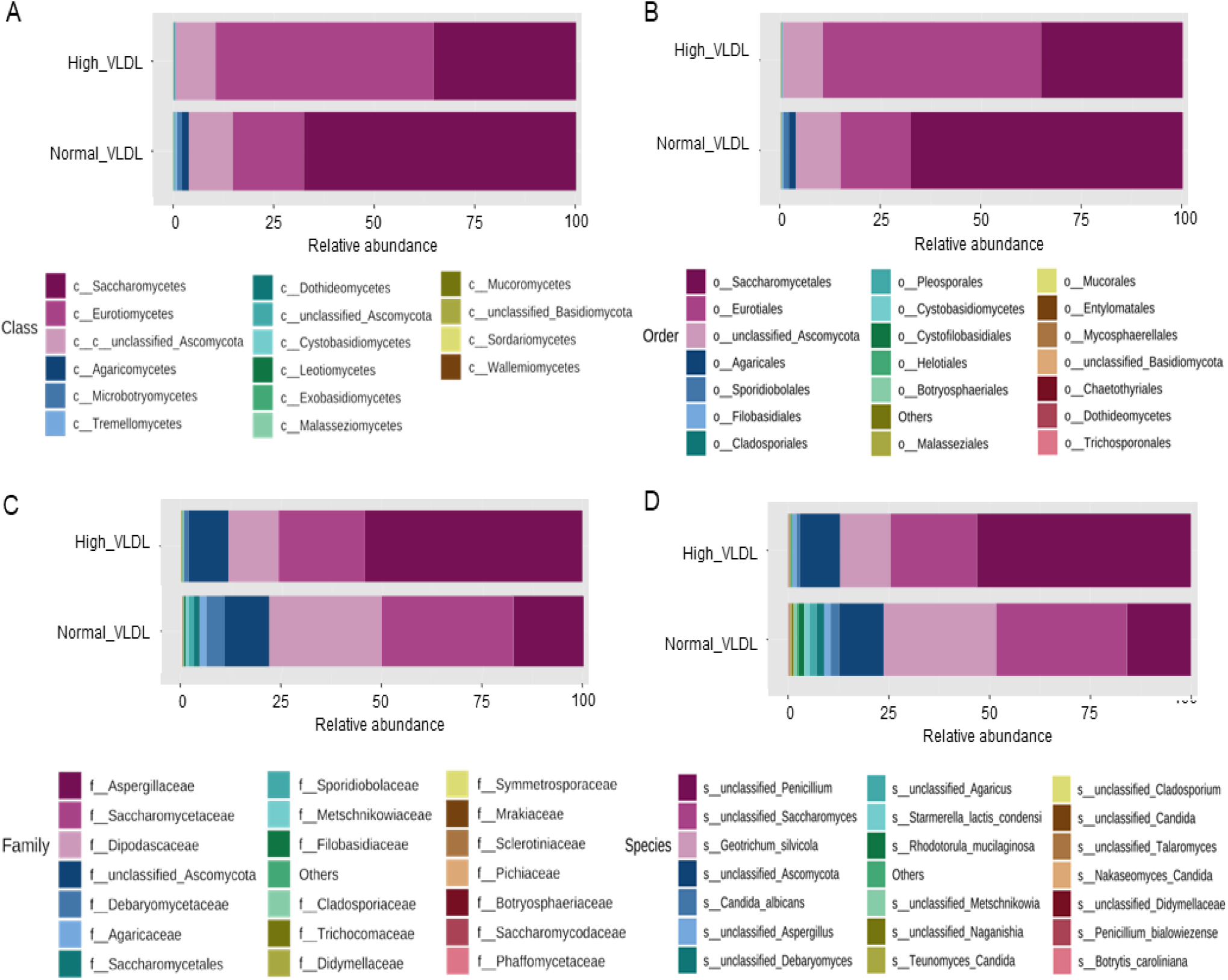
Taxonomy summary of the fungal communities associated with VLDL levels. The diagrams depicted the per-centage abundances of fungi at the (A) class; (B) order; (C) family; and (D) species levels in merged samples. Only the top 20 features are shown at the taxonomy level, with the remainder classified in Others.

**Figure S3:**
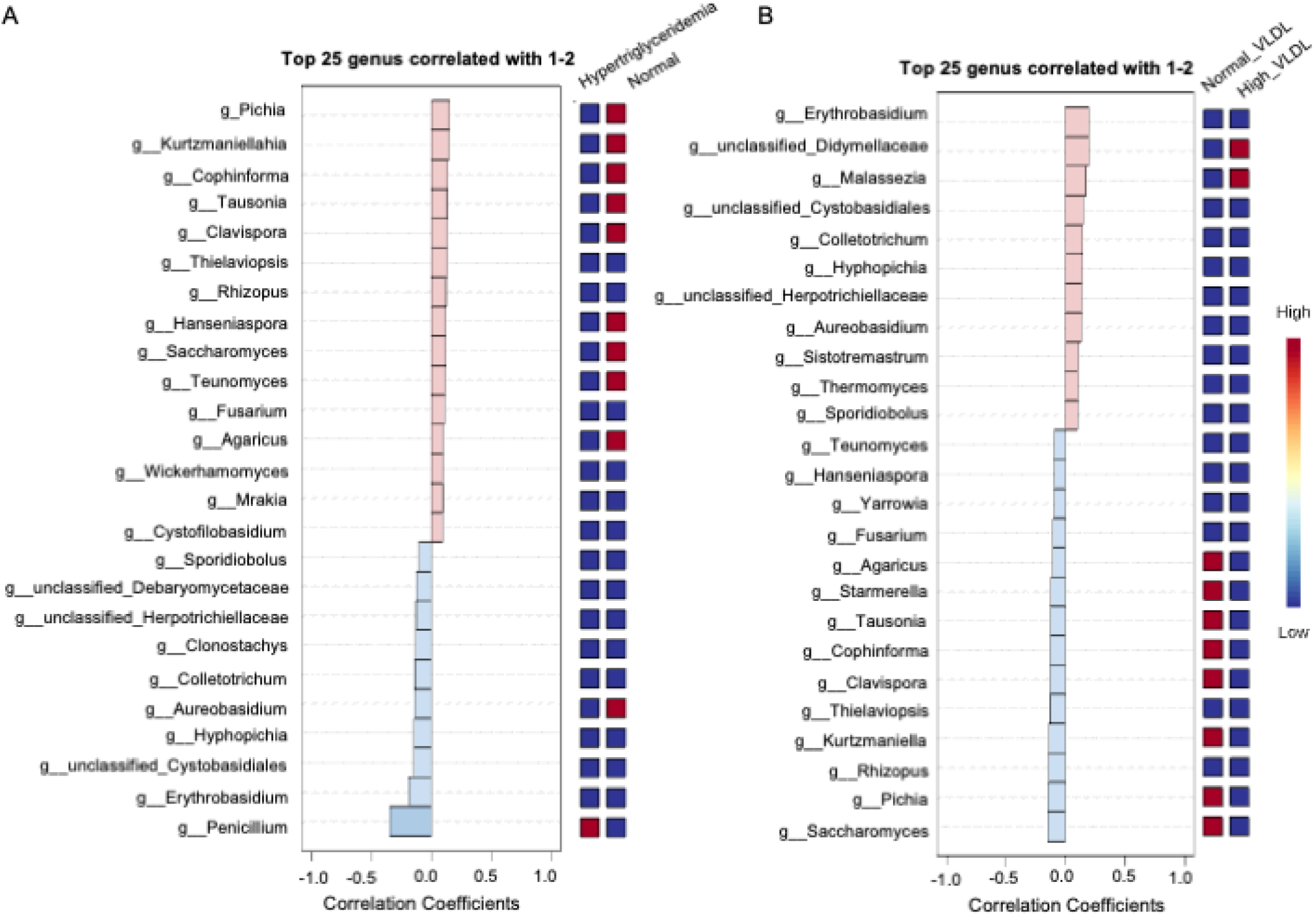
Clustering pattern of the gut mycobiome. A) The barplot illustrated the distribution pattern of identified mycobiota in the elderly Danes’ feacal samples based on the A) Tg levels; and B) VLDL levels.

**Figure S4:**
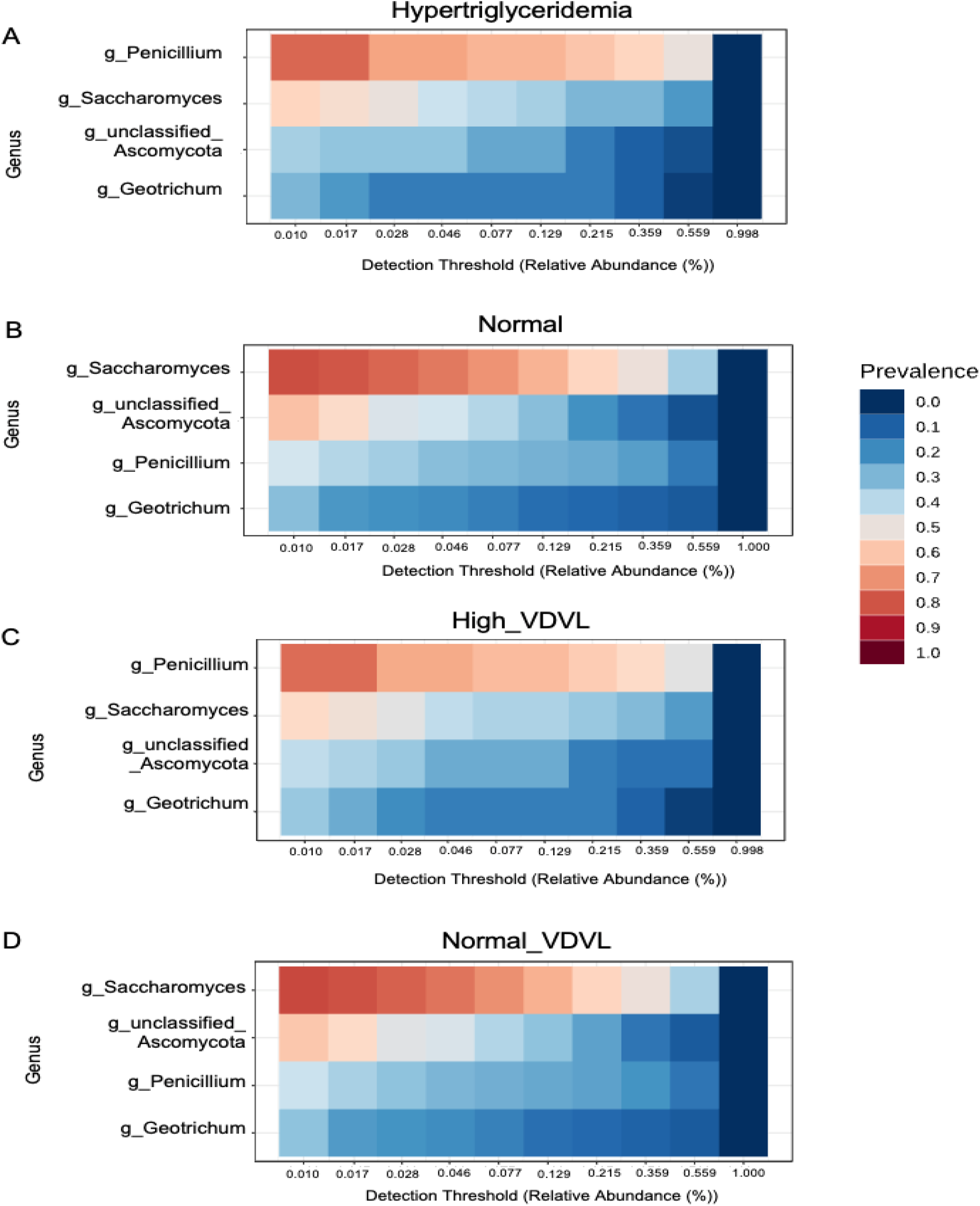
Comparison of core mycobiota at Tg and VLDL levels. The heatmaps displayed distribution of core mycobiota in sample (A) Hypertriglyceridemia; (B) Normal Tg levels; and (C) High; (D) Normal VLDL levels.

## References

[1] H. Kim et al., “Prevalence and incidence of atherosclerotic cardiovascular disease and its risk factors in Korea: a nationwide population-based study,” BMC Public Health, vol. 19, no. 1, p. 1112, 2019, doi: 10.1186/s12889-019-7439-0.

[2] T. M. Powell-Wiley et al., “Obesity and Cardiovascular Disease: A Scientific Statement From the American Heart Association,” Circulation, vol. 143, no. 21, pp. e984–e1010, May 2021, doi: 10.1161/CIR.0000000000000973.

[3] F. D. Fuchs and P. K. Whelton, “High Blood Pressure and Cardiovascular Disease,” Hypertension, vol. 75, no. 2, pp. 285–292, Feb. 2020, doi: 10.1161/HYPERTENSIONAHA.119.14240.

[4] M. Hedayatnia et al., “Dyslipidemia and cardiovascular disease risk among the MASH AD study population,” Lipids Health Dis., vol. 19, no. 1, p. 42, 2020, doi: 10.1186/s12944-020-01204-y.

[5] M. Arca et al., “Association of Hypertriglyceridemia with All-Cause Mortality and Atherosclerotic Cardiovascular Events in a Low-Risk Italian Population: The TG-REAL Retrospective Cohort Analysis,” J. Am. Heart Assoc., vol. 9, no. 19, p. e015801, Oct. 2020, doi: 10.1161/JAHA.119.015801.

[6] S. Ahmadmehrabi and W. H. W. Tang, “Gut microbiome and its role in cardiovascular diseases,” Curr. Opin. Cardiol., vol. 32, no. 6, 2017, [Online]. Available: https://journals.lww.com/co-cardiology/Fulltext/2017/11000/Gut_microbiome_and_its_role_in_cardiovascular.17.aspx.

[7] B. J. North and D. A. Sinclair, “The Intersection Between Aging and Cardiovascular Disease,” Circ. Res., vol. 110, no. 8, pp. 1097–1108, Apr. 2012, doi: 10.1161/CIRCRESAHA.111.246876.

[8] P. Alonso-Fernández and M. Fuente, “Role of the immune system in aging and longevity,” Curr Aging Sci, vol. 4, 2011, doi: 10.2174/1874609811104020078.

[9] S. Rampelli et al., “Functional metagenomic profiling of intestinal microbiome in extreme ageing,” vol. 5, no. 12, pp. 902–912, 2013.

[10] M. J. Claesson et al., “Gut microbiota composition correlates with diet and health in the elderly.,” Nature, vol. 488, no. 7410, pp. 178–84, Aug. 2012, doi: 10.1038/nature11319.

[11] S. Saraswati and R. Sitaraman, “Aging and the human gut microbiota—from correlation to causality,” Frontiers in Microbiology, vol. 5. p. 764, 2015.

[12] N. Thevaranjan et al., “Age-Associated Microbial Dysbiosis Promotes Intestinal Permeability, Systemic Inflammation, and Macrophage Dysfunction,” Cell Host Microbe, vol. 21, no. 4, pp. 455–466.e4, Apr. 2017, doi: 10.1016/j.chom.2017.03.002.

[13] A. W. Ahmad et al., “IDDF2022-ABS-0255 An observational study to identify the diversity of gut microbiome composition among obese versus healthy individuals among healthcare staffs and students in Pahang, Malaysia,” Gut, vol. 71, no. Suppl 2, p. A171 LP–A173, Sep. 2022, doi: 10.1136/gutjnl-2022-IDDF.240.

[14] S. M. Ahmad Kendong, R. A. Raja Ali, K. N. M. Nawawi, H. F. Ahmad, and N. M. Mokhtar, “Gut Dysbiosis and Intestinal Barrier Dysfunction: Potential Explanation for Early-Onset Colorectal Cancer,” Front. Cell. Infect. Microbiol., vol. 11, p. 1244, 2021, doi: 10.3389/fcimb.2021.744606.

[15] N. S. Abd Khalid et al., “IDDF2022-ABS-0263 Gut microbiome of women diagnosed with breast cancer within Pahang, Malaysia,” Gut,vol. 71, no. Suppl 2, p. A175 LP–A177, Sep. 2022, doi: 10.1136/gutjnl-2022-IDDF.244.

[16] E. Biagi et al., “Gut Microbiota and Extreme Longevity,” Curr. Biol., vol. 26, no. 11, pp. 1480–1485, Aug. 2016, doi: 10.1016/j.cub.2016.04.016.

[17] M. A. Jackson et al., “Signatures of early frailty in the gut microbiota,” Genome Med., vol. 8, no. 1, pp. 1–11, 2016, doi: 10.1186/s13073-016-0262-7.

[18] P. W. O’Toole and I. B. Jeffery, “Gut microbiota and aging,” Science (80-.)., vol. 350, no. 6265, pp. 1214–1215, Dec. 2015.

[19] S.-H. Park, K.-A. Kim, Y.-T. Ahn, J.-J. Jeong, C.-S. Huh, and D.-H. Kim, “Comparative analysis of gut microbiota in elderly people of urbanized towns and longevity villages,” BMC Microbiol., vol. 15, no. 1, pp. 1–9,2015, doi: 10.1186/s12866-015-0386-8.

[20] F. Strati et al., “Age and Gender Affect the Composition of Fungal Population of the Human Gastrointestinal Tract,” Front. Microbiol.,vol. 7, p. 1227, 2016, doi: 10.3389/fmicb.2016.01227.

[21] A. K. Nash et al., “The gut mycobiome of the Human Microbiome Project healthy cohort,” Microbiome, vol. 5, no. 1, p. 153, Nov. 2017, doi: 10.1186/s40168-017-0373-4.

[22] M. Mar Rodríguez et al., “Obesity changes the human gut mycobiome,” Sci. Rep., vol. 5, p. 14600, 2015, doi: l0.l038/srep14600/r http://www.nature.com/articles/srep14600#supplementary-information.

[23] M. Shuai et al., “Mapping the human gut mycobiome in middle-aged and elderly adults: multiomics insights and implications for host metabolic health,” Gut, vol. 71, no. 9, pp. 1812 LP–1820, Sep. 2022, doi: 10.1136/gutjnl-2021-326298.

[24] F. Zhang, D. Aschenbrenner, J. Y. Yoo, and T. Zuo, “The gut mycobiome in health, disease, and clinical applications in association with the gut bacterial microbiome assembly,” The Lancet Microbe, Sep. 2022, doi: 10.1016/S2666-5247(22)00203-8.

[25] H. F. Ahmad et al., “IDDF2020-ABS-0174 Onset of hypertriglyceridemia in relation to dietary intake, gut microbiome and metabolom-ics signatures among home dwelling elderly,” Gut, vol. 69, no. Suppl 2, p. A21 LP–A21, Nov. 2020, doi: 10.1136/gutjnl-2020-IDDF.29.

[26] C. A. Kumamoto, “The Fungal Mycobiota: Small Numbers, Large Impacts,” Cell Host Microbe, vol. 19, no. 6, pp. 750–751, Jun. 2016, doi: http://dx.doi.org/10.1016/j.chom.2016.05.018.

[27] H. Sokol et al., “Fungal microbiota dysbiosis in IBD,” Gut, Feb. 2016, doi: 10.1136/gutjnl-2015-310746.

[28] G. Liguori et al., “Fungal Dysbiosis in Mucosa-associated Microbiota of Crohn’s Disease Patients,” J. Crohn’s Colitis, vol. 10, no. 3, pp. 296–305, Mar. 2016, doi: 10.1093/ecco-jcc/jjv209.

[29] M. R. Chacón et al., “The gut mycobiome composition is linked to carotid atherosclerosis,” Benef. Microbes, vol. 9, no. 2, pp. 185–198, Nov. 2017, doi: 10.3920/BM2017.0029.

[30] J. Peng, F. Luo, G. Ruan, R. Peng, and X. Li, “Hypertriglyceridemia and atherosclerosis,” Lipids Health Dis., vol. 16, p. 233, Dec. 2017, doi: 10.1186/s12944-017-0625-0.

[31] N. Sarwar et al., “Triglycerides and the Risk of Coronary Heart Disease,” Circulation, vol. 115, no. 4, pp. 450 LP–458, Jan. 2007.

[32] N. Bg, M. Benn, P. Schnohr, and A. Tybjærg-Hansen, “Nonfasting triglycerides and risk of myocardial infarction, ischemic heart disease, and death in men and women,” JAMA, vol. 298, no. 3, pp. 299–308, Jul. 2007.

[33] Y. Yu, F. Raka, and K. Adeli, “The Role of the Gut Microbiota in Lipid and Lipoprotein Metabolism,” Journal of Clinical Medicine, vol. 8, no. 12. 2019, doi: 10.3390/jcm8122227.

[34] R. Villette et al., “Unraveling Host-Gut Microbiota Dialogue and Its Impact on Cholesterol Levels,” Frontiers in Pharmacology, vol.11. 2020, [Online]. Available: https://www.frontiersin.org/articles/10.3389/fphar.2020.00278.

[35] J. Ma and H. Li, “The Role of Gut Microbiota in Atherosclerosis and Hypertension,” Frontiers in Pharmacology,vol.9. 2018, [Online]. Available: https://www.frontiers!n.org/articles/10.3389/fphar.2018.01082.

[36] A. K. Duttaroy, “Role of Gut Microbiota and Their Metabolites on Atherosclerosis, Hypertension and Human Blood Platelet Function: A Review,” Nutrients, vol. 13, no. 1. 2021, doi: 10.3390/nu13010144.

[37] M. D. Pieczynska, Y. Yang, S. Petrykowski, O. K. Horbanczuk, A. G. Atanasov, and J. O. Horbanczuk, “Gut Microbiota and Its Metabolites in Atherosclerosis Development,” Molecules, vol. 25, no. 3. 2020, doi: 10.3390/molecules25030594.

[38] J. White, T.J., Bruns, T., Lee, S., & Taylor, “Amplification and direct sequencing of fungal ribosomal RNA genes for phylogenetics.,” in PCR Protocols: A guide to Methods and Applications., New York, USA: Academic Press, Inc, 1990, pp. 315–322.

[39] D. D. Tay, S. W. Siew, S. Shamzir Kamal, M. N. Razali, and H. F. Ahmad, “ITS1 amplicon sequencing of feline gut mycobiome of Malaysian local breeds using Nanopore Flongle,” Arch. Microbiol., vol. 204, no. 6, p. 314, 2022, doi: 10.1007/s00203-022-02929-3.

[40] C. L. Schoch et al., “Finding needles in haystacks: linking scientific names, reference specimens and molecular data for Fungi,” Database, vol. 2014, p. bau061, Jan. 2014, doi: 10.1093/database/bau061.

[41] R. L. Bechshøft et al., “Counteracting Age-related Loss of Skeletal Muscle Mass: a clinical and ethnological trial on the role of protein supplementation and training load (CALM Intervention Study): study protocol fora randomized controlled trial,” Trials, vol. 17, no. 1, p. 397, Aug. 2016, doi: 10.1186/s13063-016-1512-0.

[42] S. Chen, Y. Zhou, Y. Chen, and J. Gu, “fastp: an ultra-fast all-in-one FASTQ preprocessor,” Bioinformatics, vol. 34, no. 17, pp. i884–1890, Sep. 2018, doi: 10.1093/bioinformatics/bty560.

[43] M. Martin, “Cutadapt removes adapter sequences from high-throughput sequencing reads,” EMBnet.journal; Vol 17, No 1 Next Gener. Seg. Data Anal., 2011, doi: 10.14806/ej.17.1.200.

[44] B. J. Callahan, P. J. McMurdie, M. J. Rosen, A. W. Han, A. J. A. Johnson, and S. P. Holmes, “DADA2: High-resolution sample inference from Illumina amplicon data,” Wot. Methods, vol. 13, no. 7, pp. 581–583, 2016, doi: 10.1038/nmeth.3869.

[45] E. Bolyen et al., “Reproducible, interactive, scalable and extensible microbiome data science using QIIME 2,” Wot. Biotechnol., vol. 37, no. 8, pp. 852–857, 2019, doi: 10.1038/s41587-019-0209-9.

[46] J. Chong, P. Liu, G. Zhou, and J. Xia, “Using MicrobiomeAnalyst for comprehensive statistical, functional, and meta-analysis of microbiome data,” Wot. Protoc., vol. 15, no. 3, pp. 799–821, 2020, doi: 10.1038/s41596-019-0264-1.

[47] J. L. Castro-Mejia et al., “Physical fitness in community-dwelling older adults is linked to dietary intake, gut microbiota, and me-tabolomic signatures,” Aging Cell, vol. 19, no. 3, p. e13105, Mar. 2020, doi: https://doi.org/10.llll/acel.13105.

[48] A. Dhariwal, J. Chong, S. Habib, I. L. King, L. B. Agellon, and J. Xia, “MicrobiomeAnalyst: A web-based tool for comprehensive statistical, visual and meta-analysis of microbiome data,” Nucleic Acids Res., vol. 45, no. W1, pp. W180–W188, 2017, doi: 10.1093/nar/gkx295.

[49] J. N. Paulson, O. C. Stine, H. C. Bravo, and M. Pop, “Differential abundance analysis for microbial marker-gene surveys,” Wot. Methods,vol. 10, no. 12, pp. 1200–1202, 2013, doi: 10.1038/nmeth.2658.

[50] T. Chen, H. Zhang, Y. Liu, Y.-X. Liu, and L. Huang, “EVenn: Easy to create repeatable and editable Venn diagrams and Venn networks online,” J. Genet. Genomics, vol. 48, no. 9, pp. 863–866, Sep. 2021, doi: 10.1016/j.jgg.2021.07.007.

[51] S. W. Siew, S. M. Musa, N. ‘Azyyati Sabri, M. F. Farida Asras, and H. F. Ahmad, “The Microbiome and Metabolome Analyses of PreTreated Healthcare Wastes During Covid-19 Pandemic Reveal Potent Pathogens, Antibiotics Residues, and Antibiotic Resistance Genes Against Beta-Lactams,” SSRN Electron. J., vol. 2, no. 3, pp. 54–59, 2022, doi: 10.2139/ssrn.4240492.

[52] A. F. Members: et al.,‘ “European Guidelines on cardiovascular disease prevention in clinical practice (version 2012)’ The Fifth Joint Task Force of the European Society of Cardiology and Other Societies on Cardiovascular Disease Prevention in Clinical Practice (constituted by r,” Eur. Heart J., vol. 33, no. 17, p. 2126, Sep. 2012.

[53] Ž. Reiner, “Hypertriglyceridaemia and risk of coronary artery disease,” Wot. Rev. Cardiol., vol. 14, p. 401, Mar. 2017.

[54] G. F. Watts, E. M. M. Ooi, and D. C. Chan, “Demystifying the management of hypertriglyceridaemia,” Wot. Rev. Cardiol., vol. 10, p. 648, Sep. 2013.

[55] P. M. Hunter and R. A. Hegele, “Functional foods and dietary supplements for the management of dyslipidaemia,” Wot. Rev. Endocrinol., vol. 13, p. 278, Jan. 2017.

[56] S. M. Grundy, “Hypertriglyceridemia, insulin resistance, and the metabolic syndrome,” Am. J. Cardiol., vol. 83, no. 9, Supplement 2, pp. 25–29, 1999, doi: https://doi.org/10.1016/S0002-9149(99)00211-8.

[57] H. K. Singh, M. S. Prasad, A. K. Kandasamy, and K. Dharanipragada, “Tamoxifen-induced hypertriglyceridemia causing acute pancreatitis,” J. Pharmacol. Pharmacother., vol. 7, no. 1, pp. 38–40, Feb. 2016, doi: 10.4103/0976-500X.179365.

[58] K. P. Luna-Castillo et al., “The Effect of Dietary Interventions on Hypertriglyceridemia: From Public Health to Molecular Nutrition Evidence,” Nutrients, vol. 14, no. 5. 2022, doi: 10.3390/nu14051104.

[59] B. G. Nordestgaard and A. Varbo, “Triglycerides and cardiovascular disease,” Lancet, vol. 384, no. 9943, pp. 626–635, Aug. 2014, doi: 10.1016/S0140-6736(14)61177-6.

[60] J. Borén et al., “Low-density lipoproteins cause atherosclerotic cardiovascular disease: pathophysiological, genetic, and therapeutic insights: a consensus statement from the European Atherosclerosis Society Consensus Panel,” fur. Heart J., vol. 41, no. 24, pp. 2313–2330, Jun. 2020, doi: 10.1093/eurheartj/ehz962.

[61] A. M. Madsen et al., “Generation and Characterization of Indoor Fungal Aerosols for Inhalation Studies,” Appl. Environ. Microbiol.,vol. 82, no. 8, pp. 2479–2493, Apr. 2016, doi: 10.1128/AEM.04063-15.

[62] H. E. Hallen-Adams and M. J. Suhr, “Fungi in the healthy human gastrointestinal tract,” Virulence, vol. 8, no. 3, pp. 352–358, Apr. 2017, doi: 10.1080/21505594.2016.1247140.

[63] T. Yatsunenko, F. E. Rey, M. J. Manary, I. Trehan, M. G. Dominguez-Bello, and M. Contreras, “Human gut microbiome viewed across age and geography,” Nature, vol. 486, 2012.

[64] T. Jensen et al., “Whey protein stories - An experiment in writing a multidisciplinary biography,” Appetite, vol. 107, pp. 285–294, 2016, doi: https://doi.org/10.1016/j.appet.2016.08.010.

[65] J. Tang, I. D. Iliev, J. Brown, D. M. Underhill, and V. A. Funari, “Mycobiome: Approaches to analysis of intestinal fungi,” J. Immunol. Methods, vol. 421, pp. 112–121, Jun. 2015, doi: 10.1016/j.jim.2015.04.004.

[66] D. M. Underhill and I. D. Iliev, “The mycobiota: interactions between commensal fungi and the host immune system,” Nat Rev Immunol, vol. 14, no. 6, pp. 405–416, Jun. 2014.

[67] P. D. Scanlan and J. R. Marchesi, “Micro-eukaryotic diversity of the human distal gut microbiota: qualitative assessment using culture-dependent and -independent analysis of faeces,” ISMEJ, vol. 2, no. 12, pp. 1183–1193, Jul. 2008.

[68] A. C. Scott et al., “Chemical Mediators of the Muscle Ergoreflex in Chronic Heart Failure,” Circulation, vol. 106, no. 2, pp. 214 LP–220, Jul. 2002.

[69] L. Berglund et al., “Evaluation and Treatment of Hypertriglyceridemia: An Endocrine Society Clinical Practice Guideline,” J. Clin. Endocrinol. Metab., vol. 97, no. 9, pp. 2969–2989, Sep. 2012.

[70] T. J. Anderson et al., “2012 Update of the Canadian Cardiovascular Society Guidelines for the Diagnosis and Treatment of Dyslipidemia forthe Prevention of Cardiovascular Disease in the Adult,” Can. J. Cardiol., vol. 29, no. 2, pp. 151–167, Feb. 2013, doi: 10.1016/j.cjca.2012.11.032.

[71] T. Teramoto et al., “Executive Summary of the Japan Atherosclerosis Society (JAS) Guidelines for the Diagnosis and Prevention of Atherosclerotic Cardiovascular Diseases in Japan & 2012 Version,” J. Atheroscler. Thromb., vol. 20, no. 6, pp. 517–523, 2013, doi: 10.5551/jat.15792.

[72] Ž. Reiner et al., “ESC/EAS Guidelines forthe management of dyslipidaemiasThe Task Force forthe management of dyslipidaemias of the European Society of Cardiology (ESC) and the European Atherosclerosis Society (EAS),” Eur. Heart J., vol. 32, no. 14, pp. 1769–1818, Jul. 2011.

[73] A. L. Catapano et al., “2016 ESC/EAS Guidelines for the Management of Dyslipidaemias,” Atherosclerosis, vol. 253, pp. 281–344, Oct. 2016, doi: 10.1016/j.atherosclerosis.2016.08.018.

[74] M. Alves-Bezerra and D. E. Cohen, “Triglyceride Metabolism in the Liver,” in Comprehensive Physiology, 2017, pp. 1–22.

[75] E. Baltierra-Trejo, J. M. Sánchez-Yáñez, O. Buenrostro-Delgado, and L. Márquez-Benavides, “Production of short-chain fatty acids from the biodegradation of wheat straw lignin by Aspergillus fumigatus,” Bioresour. Technol., vol. 196, pp. 418–425, 2015, doi: https://doi.org/10.1016/j.biortech.2015.07.105.

[76] A. Risely, “Applying the core microbiome to understand host-microbe systems,” J. Anim. Ecol., vol. 89, no. 7, pp. 1549–1558, Jul. 2020, doi: https://doi.org/10.1111/1365-2656.13229.

[77] T. O. Akanbi, J. L. Adcock, and C. J. Barrow, “Selective concentration of EPA and DHA using Thermomyces lanuginosus lipase is due to fatty acid selectivity and not regioselectivity,” Food Chem., vol. 138, no. 1, pp. 615–620, 2013, doi: https://doi.org/10.1016/j.foodchem.2012.11.007.

[78] M. Aziz, F. Husson, and S. Kermasha, “Optimization of the Hydrolysis of Safflower Oil for the Production of Linoleic Acid, Used as Flavor Precursor,” Int. J. Food Sci., vol. 2015, p. 594238, 2015, doi: 10.1155/2015/594238.

[79] E. d’Avila Cavalcanti-Oliveira, P. R. da Silva, A. P. Ramos, D. A. G. Aranda, and D. M. G. Freire, “Study of Soybean Oil Hydrolysis Catalyzed by *Thermomyces lanuginosus* Lipase and Its Application to Biodiesel Production *via* Hydroesterification,” Enzyme Res., vol. 2011, p. 618692, 2011, doi: 10.4061/2011/618692.

[80] N. Matuoog, K. Li, and Y. Yan, “Immobilization of Thermomyces lanuginosus lipase on multi-walled carbon nanotubesand its application in the hydrolysis offish oil,” Mater. Res. Express, vol. 4, no. 12, p. 125402, 2017, doi: 10.1088/2053-1591/aa9d02.

[81] P. Mayser and S. Schulz, “Precipitation of free fatty acids generated by Malassezia - a possible explanation for the positive effects of lithium succinate in seborrhoeic dermatitis,” J. Eur. Acad. Dermatology Venereol., vol. 30, no. 8, pp. 1384–1389, Aug. 2016, doi: https://doi.org/10.1111/jdv.13620.

[82] S. Triana et al., “Lipid Metabolic Versatility in Malassezia spp. Yeasts Studied through Metabolic Modeling,” Frontiers in Microbiology, vol.8. 2017, [Online]. Available: https://www.frontiersin.org/articles/10.3389/fmicb.2017.01772.

[83] Y. M. DeAngelis et al., “Isolation and Expression of a Malassezia globosa Lipase Gene, LIP1,” J. Invest. Dermatol., vol. 127, no. 9, pp. 2138–2146, 2007, doi: https://doi.org/10.1038/sj.jid.5700844.

[84] C. C. Uranga, J. Beld, A. Mrse, I. Córdova-Guerrero, M. D. Burkart, and R. Hernández-Martínez, “Fatty acid esters produced by Lasiodiplodia theobromae function as growth regulators in tobacco seedlings,” Biochem. Biophys. Res. Commun., vol. 472, no. 2, pp. 339–345, 2016, doi: https://doi.org/10.1016/j.bbrc.2016.02.104.

[85] M. M. Salvatore, A. Alves, and A. Andolfi, “Secondary Metabolites of Lasiodiplodia theobromae: Distribution, Chemical Diversity, Bioactivity, and Implications of Their Occurrence,” Toxins, vol. 12, no. 7. 2020, doi: 10.3390/toxins12070457.

[86] C. C. Uranga, J. Beld, A. Mrse, I. Córdova-Guerrero, M. D. Burkart, and R. Hernández-Martínez, “Data from mass spectrometry, NMR spectra, GC-MS of fatty acid esters produced by Lasiodiplodia theobromae,” Data Br., vol. 8, pp. 31–39, 2016, doi: https://doi.Org/10.1016/j.dib.2016.05.003.

[87] S. Raimondi et al., “Longitudinal Survey of Fungi in the Human Gut: ITS Profiling, Phenotyping, and Colonization,” Frontiers in Microbiology, vol.10. 2019, [Online]. Available: https://www.frontiersin.org/articles/10.3389/fmicb.2019.01575.

[88] J. A. Takahashi, B. V. R. Barbosa, B. de A. Martins, C. P. Guirlanda, and M. A. F. Moura, “Use of the Versatility of Fungal Metabolism to Meet Modern Demands for Healthy Aging, Functional Foods, and Sustainability,” Journal of Fungi, vol. 6, no. 4. 2020, doi: 10.3390/jof6040223.

[89] B. Sandstrom, N. Lyhne, J. I. Pedersen, A. Aro, I. Thorsdottir, and W. Becker, Nordic nutrition: Recommendations 2012, vol. 40, no. 4. 2012.

[90] S. J. Ott et al., “Fungi and inflammatory bowel diseases: alterations of composition and diversity,” Scand J Gastroenterol, vol. 43, 2008, doi: 10.1080/00365520801935434.

[91] A. Hot et al., “Fungal Internal Carotid Artery Aneurysms: Successful Embolization of an Aspergillus-Associated Case and Review,” Clin. Infect. Dis., vol. 45, no. 12, pp. e156–e161, Dec. 2007, doi: 10.1086/523005.

[92] S. Rønnow Schacht et al., “Investigating Risk of Suboptima I Macro and Micronutrient Intake and Their Determinants in Older Danish Adults with Specific Focus on Protein Intake—A Cross-Sectional Study,” Nutrients, vol. 11, no. 4. 2019, doi: 10.3390/nu11040795.

[93] T. A. Auchtung et al., “Investigating Colonization of the Healthy Adult Gastrointestinal Tract by Fungi,” mSphere, vol. 3, no. 2, pp. e00092–18, Mar. 2018, doi: 10.1128/mSphere.00092-18.

[94] G. Dubois, C. Girard, F.-J. Lapointe, and B. J. Shapiro, “The Inuit gut microbiome is dynamic over time and shaped by traditional foods,” Microbiome, vol. 5, no. 1, p. 151, Nov. 2017, doi: 10.1186/s40168-017-0370-7.

[95] H. Bartolomaeus et al., “Short-Chain Fatty Acid Propionate Protects From Hypertensive Cardiovascular Damage,” Circulation, vol. 139, no. 11, pp. 1407–1421, Mar. 2019, doi: 10.1161/CIRCULATIONAHA.118.036652.

[96] J. Li et al., “The fungal community and its interaction with the concentration of short-chain fatty acids in the faeces of Chenghua, Yorkshire and Tibetan pigs,” Microb. Biotechnol., vol. 13, no. 2, pp. 509–521, Mar. 2020, doi: https://doi.org/10.1111/1751-7915.13507.

[97] D. Maciejewska et al., “The short chain fatty acids and lipopolysaccharides status in sprague-dawley rats fed with high-fat and high-cholesterol diet,” J. Physiol. Pharmacol., vol. 69, no. 2, pp. 205–210, 2018, doi: 10.26402/jpp.2018.2.05.

[98] S. Deleu, K. Machiels, J. Raes, K. Verbeke, and S. Vermeire, “Short chain fatty acids and its producing organisms: An overlooked therapy for IBD?,” eBioMedicine, vol. 66, Apr. 2021, doi: 10.1016/j.ebiom.2021.103293.

